# A *de novo* missense mutation in *TUBA1A* results in reduced neural progenitor survival and differentiation

**DOI:** 10.1101/201814

**Authors:** Ashley M. Driver, Amy L. Pitstick, Chris N. Mayhew, Beth Kline-Fath, Howard M. Saal, Rolf W. Stottmann

## Abstract

Mutations in tubulins have been implicated in numerous human neurobiological disorders collectively known as “tubulinopathies.” We identified a patient with severe cortical dysgenesis and a novel *de novo* heterozygous missense mutation in *Tubulin Alpha 1a* (*TUBA1A*, c.1225 G>T). Induced pluripotent stem cells derived from this individual were differentiated into two dimensional neural rosette clusters to identify underlying mechanisms for the severe cortical dysgenesis phenotype. Patient-derived clones showed evidence of impaired neural progenitor survival and differentiation with abnormal neural rosette formation, increases in cell death, and fewer post-mitotic neurons. These features correlate with the drastically underdeveloped cortical tissues seen in the proband. This is the first experimental evidence in human tissue suggesting a mechanism underlying the role for *TUBA1A* in cortical development.

**SUMMARY STATEMENT:** Variants in tubulin genes often lead to severe congenital brain malformations. Here we identify a new mutation in *TUBA1A* and use iPSCS to show this alters proliferation, differentiation and survival of neural progenitors.

## Introduction

Development of the central nervous system (CNS) depends on a tightly regulated series of events involving cellular proliferation, migration, and differentiation. Neural progenitors must successfully undergo these events to form mature post-mitotic neurons necessary for cortical expansion. Microtubules have critical roles in these functions including regulating cell shape, division, and migration (Ganguly et al., 2012; Prosser and Pelletier, 2017). Neurons are especially dependent on microtubules as post-mitotic neurons rely on these structures to form proper neurite extensions and conduct intracellular trafficking (Dent and Baas, 2014). Additionally, microtubules have been implicated to have an essential role in establishing neuronal polarity (Conde and Caceres, 2009). Microtubules are cytoskeletal polymers consisting of linked heterodimer subunits (Conde and Caceres, 2009). These subunits consist of dimeric proteins known as α-and β-tubulins (Conde and Caceres, 2009).

The “tubulinopathies” are a subset of complex cortical malformations specifically caused by mutations in tubulin genes. Mutations in both α-and β-tubulin genes have been reported in patients with cortical abnormalities including microcephaly, lissencephaly, and cognitive deficits (Bamba et al., 2016; Romaniello et al., 2015; Tischfield et al., 2011). There are over 45 known *TUBA1A* mutations that have been identified in patients with a spectrum of CNS defects (Romaniello et al., 2015; Shimojima et al., 2014). However, the specific cellular defects that underlie these CNS phenotypes are poorly defined.

We utilized whole exome sequencing to identify a novel *de novo* heterozygous missense *TUBA1A* mutation in a patient with severe cortical dysgenesis. Additionally, we use patient-derived induced pluripotent stem cells (iPSCs) to address potential mechanisms *in vitro* and identify numerous cellular phenotypes that can together likely explain the disrupted cortical development seen in the proband. This study adds to the growing database of tubulin mutations and, for the first time, identifies potential cellular defects that contribute to CNS phenotypes in affected patients.

## Results

### Patient features

The patient was the male product of a vaginal delivery at 37 6/7 weeks to a 32 year old G4P201 woman and her 34 year old unrelated husband. The pregnancy was complicated by identification of multiple anomalies noted on high resolution ultrasound, including hydrocephalus, abnormal brainstem and hypoplastic cerebellum with absent vermis, and a large interhemispheric cyst with abnormal sulcation. There were also bilateral clubfeet. The patient was seen in the Fetal Care Center and a fetal MRI scan was performed at 22 3/7 weeks confirmed the ultrasound findings and identified additional findings including aqueduct stenosis, macrocephaly, and several additional findings. Non-invasive prenatal testing with free fetal DNA showed that there was no aneuploidy and a fetal echocardiogram was normal. The family decided to continue pregnancy and decided that given the poor prognosis to offer palliative care at delivery. The infant was delivered via Cesarean section because of cephalopelvic disproportion. Apgar scores were 2 and 2 at one and five minutes, respectively. Birth weight and length were not obtained. Comfort care was provided and the infant expired at about 8 hours of age. Parents declined autopsy, but permitted post-mortem MRI and genomic testing.

Examination of the infant post mortem showed a normal sized fetus with enlarged head. The anterior fontanelle was 4 cm. There was frontal bossing with high and wide forehead, the eyes were deeply set and there was a depressed nasal bridge, square faced, mild micrognathia. The nose was short, philtrum was formed and short and mouth was small and the upper and posterior vermilions were thin. The cheeks were full and the palate was intact. The ears were normally placed and there was overfolding of the superior helix. Chest was symmetric. There was campodactyly of the fingers with no other digital anomalies. Lower extremities were mildly bowed, there was bilateral talipes equinovarus with metatarsus adductus. No additional skeletal anomalies were noted. Genitalia were normal male.

### Family History

The family history is positive for the patient’s mother having a balanced translocation between the long arms of chromosomes 11 and 22, 46,XX,t(11;22)(q23;q11.2). His mother’s first pregnancy resulted in a stillbirth male at 37 weeks gestation, presumably from a cord accident. No ultrasound anomalies had been identified, no autopsy or genetic studies were performed. The second pregnancy was a first trimester miscarriage and the third pregnancy resulted in a girl who has Noonan syndrome and a *PTPN11* de novo pathogenic variant. There is a distant paternal cousin with mild autism spectrum disorder and a maternal great uncle who was diagnosed with phenylketonuria at 2 years and has severe developmental disability.

### MRI Imaging

A post-mortem MRI revealed severe lateral ventriculomegaly with bilateral ventricular rupture and absence of the septum pellucidum (Fig. 1C). There was moderate third ventriculomegaly with absence of CSF in the aqueduct of Slyvius in association with a thickened tectum, consistent with aqueduct obstruction (Fig. 1D). There was no evidence of heterotopias along the ventricular surface, although cysts were noted at the caudothalamic ganglionic eminences (Fig. 1C). The supratentorial brain mantle was reduced in size and there was an overall significant loss in white matter volume. The sulcation was shallow without evidence for cobblestone lissencephaly (Fig. 1C). The corpus callosum was nearly absent with thin remnant in the area of the genu. The hippocampal tissue was thin and the thalamus appeared partially fused. The basal ganglia structures were poorly defined with lack of visualization of the internal capsule (Fig. 2E). The brainstem was small with a “Z” shaped configuration (Fig. 2D) and small central cleft. The cerebellar hemispheres and vermis were small with limited sulcation. The optic nerves were hypoplastic and olfactorary bulbs absent. Outside the CNS, we observed mild micrognathia but no evidence of cleft lip/palate.

**Figure 1.**
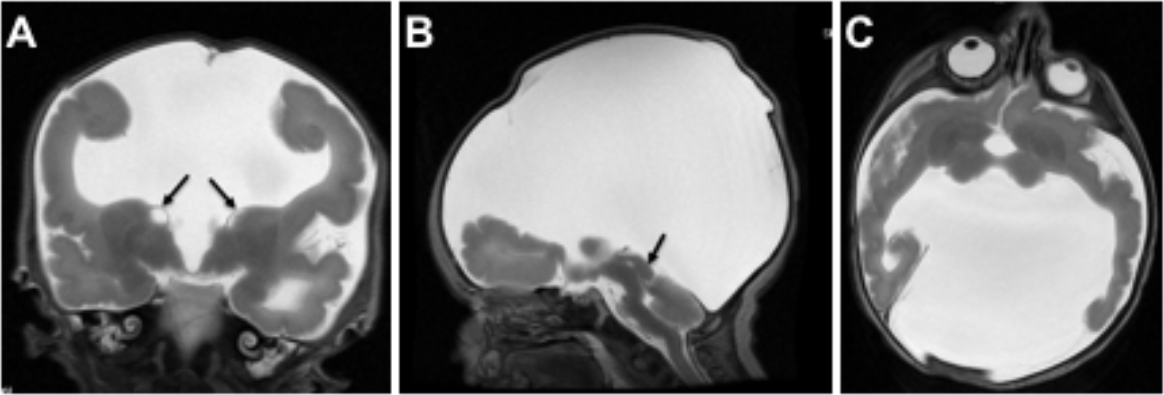
(A-C) MRI Imaging. (A) Coronal T2-weighted MRI at 38 weeks of gestation showing severe lateral ventriculomegly and moderate third ventriculomgely with absence of the septum pellucidum. The brain mantle is thin with shallow sulcation. Small cysts are present at the ganglionic eminence present at the caudothalamic groove (arrows). (B) Sagittal T2 image demonstrates lack of CSF distention of the aqueduct of Slyvius with thickened tectum (arrow). The brainstem is small and demonstrates a “Z” shaped configuration. (C) The basal ganglia structures are small with lack of visualization of normal internal capsule.

**Figure 2.**
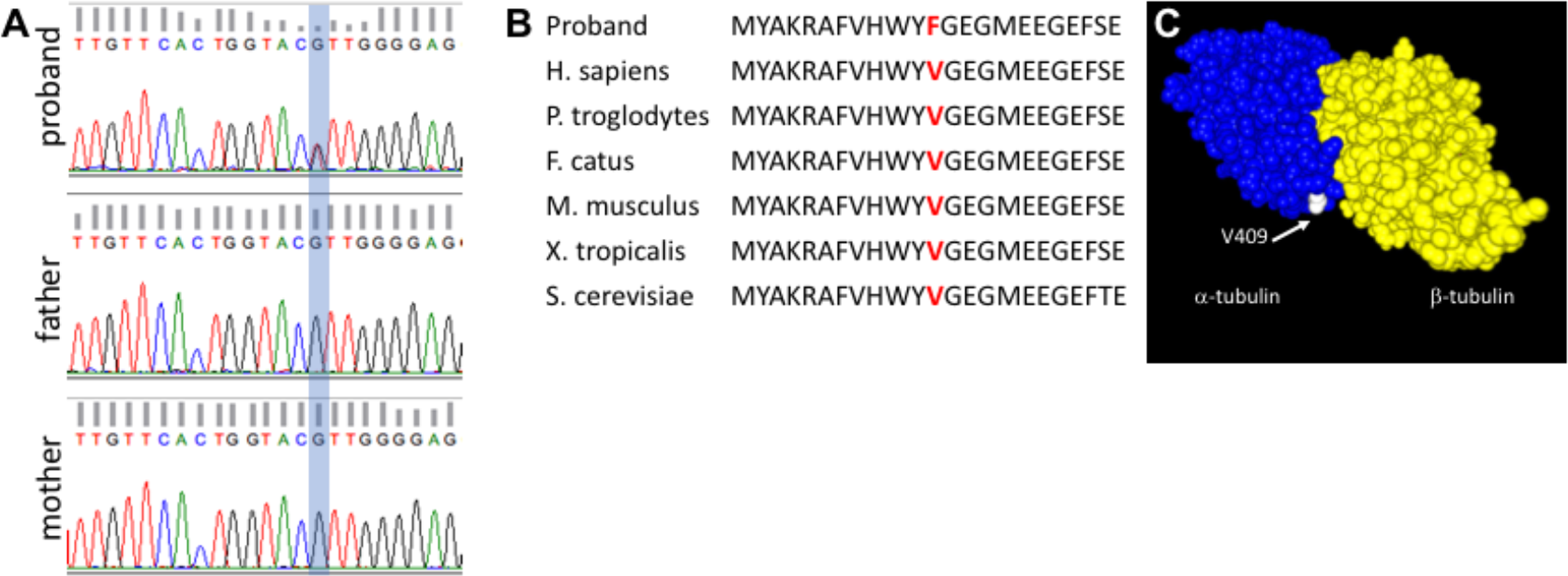
(A) Sanger sequencing validation of the exome sequencing results. Blue shading highlights the heterozygous *TUBA1A* variant. (B) Protein alignment shows the conserved nature of the amino acid residue altered in the p.V409F patient. (C) Three-dimensional modeling of the variant highlights the exterior position of the V409 amino acid (white) on the α-tubulin protein.

### Exome results identify a de novo missense variant in TUBA1A

Chromosome analysis for the infant showed that he had a maternally inherited balanced translocation, 46,XY,t(11;22)(q23;q11.2)mat. This is a known translocation and is not unique to the proband in this pedigree (Zackai and Emanuel, 1980) so we therefore conclude this is not related to the CNS malformations. Whole exome sequencing (Illumina Hi Seq 2000) was performed for the affected child and parents after obtaining informed consent according to our Cincinnati Children’s Hospital Medical Center (CCHMC) IRB-approved protocol. Alignment and variant detection was performed using the Broad Institute’s web-based Genome Analysis Toolkit (GATK; Genome Reference Consortium Build 37) (McKenna et al., 2010). Quality control and data filtering were performed on VCF files independently using manual curation and Golden Helix’s SNP and Variation Suite. The initial bioinformatics analysis identified 144,624 variants (Table 1). After filtering for quality control, coding variants, and minor allele frequencies (variants present in dbSNP) we tested three inheritance models. No compound heterozygous mutations were found and only one recessive mutation was identified. The recessive variant is in a gene with known roles in dentin biology *(DSPP*), so we did not consider it causal in this family. Analysis for *de novo* variants identified 7 candidates. Two were not supported by a new analysis of the bam files and considered technical artifacts and another was seen 14 times in an internal cohort and was thus considered a geographically localized variant or a technical artifact in our sequencing apparatus. Three more variants were in genes with known mutations in mouse or human which were not consistent with the proband (*LRGUK*, *NLRP1*, *LILRB3*). The final variant was a missense variant in *TUBULIN, ALPHA-1A (TUBA1A):* c.1225G>T (p.V409F). This variant was confirmed by Sanger sequencing variant and was not present in either parent (Fig. 2A). Furthermore, this variant was not identified in the Exome Aggregation Consortium (ExAC) database or 1000 genomes. The c.1225 G>T variant was predicted as “disease causing” by MutationTaster and “possibly damaging” by PolyPhen-2 (score: 0.932; sensitivity: 0.80; specificity: 0.94) and results in an amino acid change from a Valine to a Phenylalanine at a conserved residue (Fig. 2B). The affected residue is predicted to be on the exterior of the a-tubulin monomer, near the adjacent β-monomer (Fig. 2C). Thus, we predict the effect to be on binding of other molecules such as microtubule associated proteins or motor proteins like kinesin, rather than tubulin monomer interactions.

**Table 1.** Exome Variant Analysis.

| Familial Variant Analysis | # of Variants |
| --- | --- |
| Total Variants | 144,624 |
| Quality Control (GQ≥90) | 77,265 |
| Coding variants | 14,890 |
| Not in dbSNP | 531 |
| *Compound heterozygous* | 0 |
| *Recessive inheritance* | 1* |
| *De novo* | 7 |
| Not present in internal control | 6 |
| Supported by bam file analysis | 4 |
| Biological plausibility | 1 |

### Patient-derived iPSCs show abnormal proliferation, differentiation, and increased cell death during neural differentiation

In order to begin to probe cellular defects that may contribute to abnormal CNS development in individuals harboring the c.1225 G>T *TUBA1A* mutation, we generated patient-specific iPSCs. To serve as controls, iPSCs were also generated from fibroblasts isolated from an unrelated healthy individual. Fibroblasts were reprogrammed using episomal plasmids resulting in two clonal lines derived from the c.1225 G>T *TUBA1A* patient (241.12 and 241.15), and one clonal line from the control (115.1). The c.1225 G>T variant was present in patient derived lines and absent in the control iPSC line (Fig. 3A-C). All iPSC lines showed stereotypical pluripotent stem cell morphology with compact flattened colonies (Fig. 3D-F), consisting of cells with a high nuclear to cytoplasmic ratio (Robinton and Daley, 2012). Immunocytochemistry for SOX2, a key pluripotency factor, demonstrated uniform expression in all undifferentiated iPSCs (Fig. 3G-I). Finally, karyotype analysis demonstrated that whilst 115.1 controls exhibited a normal male karyotype, both 241.12 and 241.15 harbored the balanced translocation identified in the patients prior to the iPSC generation. Together, these data demonstrate pluripotency marker expression, retention of the predicted karyotype and preservation of the c.1225 G>T *TUBA1A* sequence in iPSCs derived from the patient with severe cortical dysgenesis.

**Figure 3.**
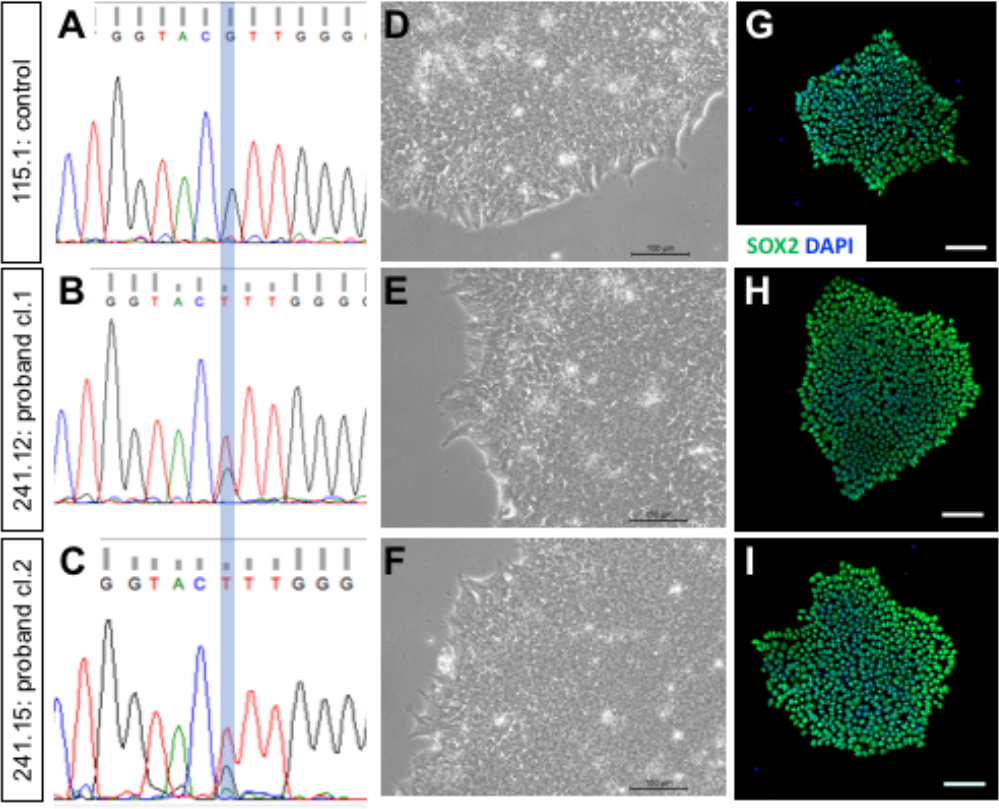
(A-C) Sanger sequencing confirmed the presence of the *TUBA1A* heterozygous c.1225 G>T mutation in patient clones and absence in control. (D-F) Patient iPSCs (241.12, 241.15) show similar colony and cell morphology between control and proband-derived iPSCs (115.1). (G-I) Immunocytochemistry for SOX2 is similar across all cells lines in this study. Scale bars in phase images are 100uM.

To begin to determine an underlying molecular mechanism(s) for the cortical malformations, iPSCs were differentiated to neural progenitors following a protocol designed to generate two dimensional rosettes. At 7 and 11 days *in vitro* (DIV), these conditions facilitate formation of morphologically identifiable rosette structures with apical-basal polarity within differentiating colonies. While the control lines had rosettes throughout the cell clusters, both patient lines showed significantly fewer organized rosettes (Fig. 4A-F). This was further confirmed with nuclear DAPI staining, which showed large circular rosettes in the control compared to very small circular structures and a significant lack of rosette formation in the patient-derived cells. Immunocytochemical analysis using PAX6 to identify neural progenitors confirmed that while the patient cultures contained progenitors, they were less radially organized. In addition, immunocytochemistry for pHH3-postive proliferative cells showed mitotically active cells along the apical surface of the control rosettes. Proliferative cells in the patient lines appeared relatively disorganized (Fig 4H-L-I). Furthermore, we detected increased numbers of Cleaved Caspase 3-positive (CC3) cells throughout patient-derived clusters as compared to control iPSCs, suggesting marked cell death is occurring during neural induction (Fig. 4M-O). Finally, analysis of beta-III tubulin, a marker for post-mitotic, differentiated neurons, revealed reduced neuronal differentiation in the patient clones as compared to control at 11 DIV (Fig. 4P-R). Taken together, these results indicate abnormal neural progenitor differentiation from patient-derived iPSCs and suggest the cortical malformations resulting from the missense mutation in *TUBA1A* are due to a combination of disorganized patterns of cellular proliferation, increased cell death and reduced differentiation. To our knowledge, this is the first study suggesting a cellular mechanism for the tubulinopathy-related cortical malformations.

**Figure 4.**
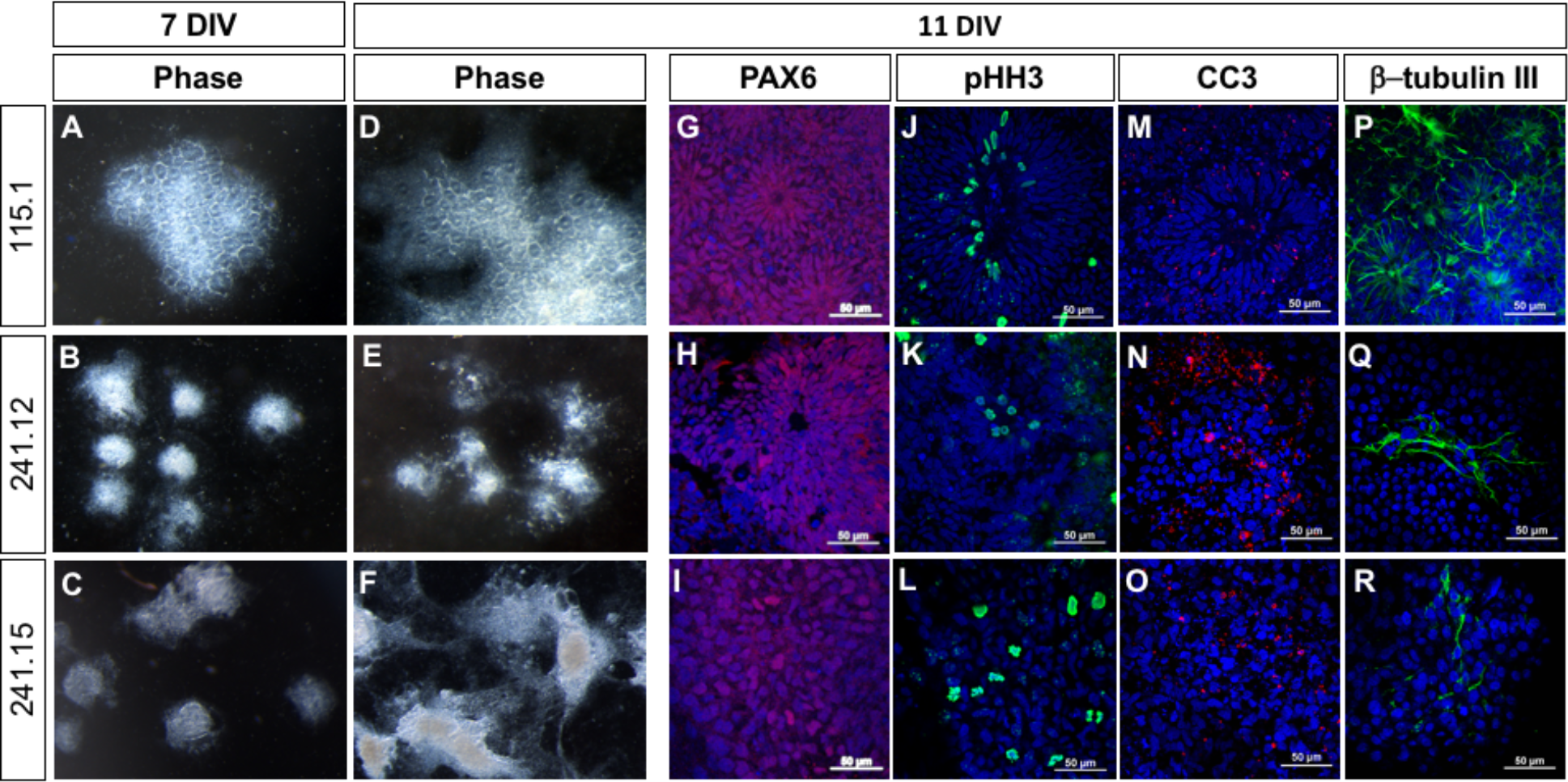
(A,D) Neural rosettes are present throughout control cell clusters at both 7 and 11 DIV. (B,C,E,F) Both patient-derived lines show abnormal morphology at. (G-I) PAX6 immuno-positive neural progenitors (red) organize into rosette structures in the control cells (G) and have lost this organization in the patient-derived lines (H,I). (J-L) pHH3 staining shows mitotic cells along the apical surface of control rosettes (J) while proliferation appears less organized in the patient-derived lines (K,L). (M-O) Immunostaining for cleaved caspase 3 (CC3) to highlight cells undergoing cell death indicates a marked increase in patient-derived lines (N,O) compared to control (M). (P-R) Immunostaining for beta-III tubulin to mark post-mitotic neurons showed decreased differentiated neurons in the patient-derived cultures (Q,R) compared to control (P). Scale bars in G-R are 50μm.

## Discussion

In this study, we identify a *de novo* heterozygous missense *TUBA1A* mutation in a patient with severe cortical malformations. We have used patient-derived cells to model brain development *in vitro* to begin to study the molecular mechanism underlying the tubulinopathies. These experiments allow us to show multiple cellular effects which we link to the *TUBA1A* variant. This include reduced and disorganized proliferation, decrease differentiation and increases in apoptosis. To our knowledge, this is the first study to suggest a mechanism for the drastic effects seen upon genetic perturbation of the tubulin loci.

While the phenotypes we observed in the proband are within the greater tubulinopathy spectrum, they are among the most severe reported for a tubulin mutation. *TUBA1A* has been previously associated with neuronal migration defects and lissencephaly; however, these phenotypes were not observed in our patient. Recent findings suggest the spectrum of *TUBA1A*-related brain malformations may be broader than initially thought (Yokoi et al., 2015). Indeed, the phenotypes observed in this study are some of the most severe seen in *TUBA1A* patients (Fallet-Bianco et al., 2014; Yokoi et al., 2015). Shared features include an absence of a septum pellucidum, poor formation of basal ganglia, a significantly small/dysmorphic brain stem, and evidence of hydranencephaly (Yokoi et al., 2015). The sheer extent of cortical tissue loss is unique, as is the aqueductal stenosis. He was also found to have a balanced maternally inherited chromosome translociation between the long arms of chomosomes 11 and 22. This is the most common chromosome translocation and it has no phenotypic manifestations (Zackai and Emanuel, 1980).

Recent studies have identified other patients with missense mutations in *TUBA1A* at the same protein residue (p.V409A and p.V409I-(Bahi-Buisson et al., 2014); see Table 2). In the p.V409I patient, central pachygyria was the primary phenotype observed without any further structural deficit. In contrast, both the p.V409A (Bahi-Buisson et al., 2014) and p.V409F (this study) show severe loss of cortical tissue formation, with the proband in this study being the most severe. Of the three missense mutations, the p.V409I is the most conservative amino acid change with the isoleucine residue maintaining a hydrophobic side chain. The p.V409A variant substitutes the small alanine residue for the hydrophobic valine. In contrast, the p.V409F variant introduces a large phenylalanine aromatic side-chain. Given the extreme phenotypic consequences of this mutation, we suspect this disrupts more of the functions of the tubulin monomer, perhaps explaining the more significant effect on neurodevelopment in the proband.

**Table 2.** Comparison of effects of pV409 variants.

|  | **Proband** | **LIS_TUB_011_**<br>**fœtus23** | **LIS_TUB_032** |
| --- | --- | --- | --- |
| **Nucleotide** | c.1225 G > T | c.1226 T > C | c.1225G > A |
| **Protein** | p.V409F | p.V409A | p.V409I |
| **Inheritance** | *de novo* | *de novo* | *de novo* |
| **Cortical Dysgenesis** | Moderate ventriculomegaly, significant loss of white matter volume, shallow sulcation without evidence of cobblestone lissencephaly | Lissencephaly with cerebellar hypoplasia | Central pachygyria |
| **Corpus Callosum** | Nearly completely absent | Complete agenesis | Dysmorphic and hypoplastic |
| **Cerebellum** | Severe vermian hypoplasia with limited sulcation | Severe vermian hypoplasia | normal |
| **Age** | 37 GW | 32 GW | 10y |
| **Basal ganglia** | Poorly defined with no visualization of internal capsule | dysmorphic | Not commented on |
| **Other notes** | Aqueduct obstruction, hypoplastic optic nerve, absent olfactory bulbs, mild micrognathia, no evidence of cleft palate |  |  |
| **Reference** | This paper | (Bahi-Buisson et al., 2014)(Fallet-Bianco et al., 2014) | (Bahi-Buisson et al., 2014) |

There is increasing evidence that tubulins may play a significant role in the survival and differentiation of developing neural cells and this is likely a causal mechanism of many tubulinopathies. In a mouse model with a homozygous mutation in *Tubb2b*, cortical development is severely perturbed due to abnormal proliferation and increased apoptosis (*Tubb2b*^*brdp/brdp*^;(Stottmann et al., 2013). This study also reported a reduction in the number of post-mitotic neurons in the *Tubb2b*^*brdp/brdp*^ brains (Stottmann et al., 2013). Similarly, recent findings on the β-tubulin gene *TUBB5* show that alterations in mitotic spindle orientation can also result in a shift from cell survival to cell death (Breuss et al., 2016). Taken together, reduced cell survival and neuron production seem to be common features of tubulin mutations leading to cortical malformations of development.

As few mouse models exist for tubulin mutations, this study highlights the potential utility of the hPSCs to model early stages of neural development and disease. As shown here, the hPSCs can be directed to a neural fate and used to form two-dimensional rosettes which recapitulate many aspects of early neural development. By studying the cellular mechanisms of development in the rosette, we can understand some of the underlying pathology in the human malformation. Here we demonstrated multiple stages of development are perturbed in the patient-derived cultures: proliferation, death and differentiation. Future efforts can include further development of the cells into three-dimensional structures to more completely model the phenotype, and/or the creation of mouse models to specifically mimic variants seen in human. In conclusion, our study identifies a new mutation in *TUBA1A* and provides evidence in patient-derived iPSCs for alpha tubulins to play a significant role in neural progenitor cell survival and differentiation.

## Materials and Methods

### Patient sequencing and variant confirmation

Informed consent was obtained according to Cincinnati Children’s Hospital Medical Center (CCHMC) institutional review board protocol #2014-3789. Following consent, whole blood was collected on the parents, residual DNA and fibroblasts from the proband. Library generation, exome enrichment, sequencing, alignment and variant detection were performed in the CCHMC Genetic Variation and Gene Discovery Core Facility (Cincinnati, OH). Briefly, sheared genomic DNA was enriched with NimbleGen EZ Exome V2 kit (Madison, WI). The exome library was sequenced using Illumina’s Hi Seq 2000 (San Diego, CA). Alignment and variant detection was performed using the Broad Institute’s web-based Genome Analysis Tookit (GATK)(McKenna et al., 2010). All analyses were performed using Genome Reference Consortium Build 37.

### Variant Filtering and Pathogenicity Assessment

Quality control and data filtering were performed on VCF files in Golden Helix’s SNP and Variation Suite (Bozeman, MT). Non-synonymous coding variants were compared to three control databases, including NHLBI’s ESP6500 exome data (Fu et al., 2013), the 1000 genomes project (Genomes Project et al., 2010), EXAC Browser (Karczewski et al., 2017) and an internal CCHMC control cohort (Patel et al., 2014). Remaining variants were subject to autosomal recessive analysis with emphasis on homozygous recessive variants found in the region of homozygosity identified by SNP microarray. The identified variant was compared to known disease genes in the OMIM and Human Gene Mutation (HGMD) (Stenson et al., 2014) databases, and to reported variants in dbSNP (Sherry et al., 2001) and the Exome Variant Server. The variant was also analyzed using Interactive Biosoftware’s Alamut v2.2 (San Diego, CA) to determine location of mutation within a protein domain, the conservation of the amino acid, the Grantham score (Grantham, 1974) and the designation of the mutation by three existing in-silico software tools, SIFT (Li et al., 2012), Polyphen (Adzhubei et al., 2010) and Mutation Taster (Schwarz et al., 2010).

### iPSC generation and maintenance

Fibroblasts were collected from the proband under an IRB-approved protocol and reprogrammed into induced pluripotent stem cells by transfection of episomal plasmids expressing Oct3/4, Sox2, Klf4, LMyc, Lin28, and shRNA against p53 as previously described (McCracken et al., 2014; Okita and Yamanaka, 2011). Established lines were maintained in a feeder-free environment using mTesR1 media (StemCell Technologies) on Matrigel (Corning) coated culture plates. Cells were maintained in a water-jacked incubator at 37°C with 5% CO_2_ and were passaged every 4-6 days using either Dispase (StemCell Technologies) or Gentle Cell Dissociation Reagent (StemCell Technologies). Standard metaphase spreads and G-banded karyotypes were determined by the CCHMC Cytogenetics Laboratory.

### 2-D neural rosette differentiation

Both control and patient-derived iPSCs were differentiated into 2-D neural rosette cultures using the StemDiff Neural Induction Protocol (StemCell Technologies). Briefly, iPSC colonies were dissociated into a single cell suspension and plated into an Aggrewell 800 plate (StemCell Technologies) for embryoid body (EB) formation in Neural Induction Media (NIM; StemCell Technologies). After 5 days EBs were re-plated on to Matrigel-coated (Corning) glass coverslips in a 24-well culture plate. Cells were then maintained in NIM until Day 7 or 11 DIV when cells were collected for immunostaining.

### Immunocytochemistry

Cells were washed with phosphate buffered saline (PBS) and fixed in 4% paraformaldehyde (PFA) for 10 minutes. For iPSC staining, cells were incubated in primary antibody (Sox2, 1:500, Santa Cruz, sc17320) diluted in blocking buffer (4% goat serum in PBS plus Triton 100X). For neural rosette staining primary antibodies for Pax6 (1:500, Abcam, ab5790), TuJ1 (1:500, SIGMA, clone 2G10), CC3 (1:300, Cell Signaling, #9961), and pHH3 (1:500, Sigma) were diluted in blocking buffer. Cells were incubated in diluted primary antibody at 4-degrees overnight followed by washes in PBS. Cells were then incubated in secondary antibody (Alexa Fluor 488/594, 1:500, Life Technologies) diluted in goat serum plus PBS-T for 1 hour at room temperature in the dark. Following incubation, cells were washed with PBS, incubated with DAPI (1:1000) and washed in PBS. Coverslips were sealed with ProLong Gold Anti-fade Reagent (Life Technologies). All images were taken using a Nikon C2 Confocal microscope. All paired images were taken at the same magnification.

